# Spatial Variation of Microtubule Depolymerization in Large Asters Suggests Regulation by MAP Depletion

**DOI:** 10.1101/2020.06.26.172783

**Authors:** Keisuke Ishihara, Franziska Decker, Paulo Caldas, James F. Pelletier, Martin Loose, Jan Brugués, Timothy J. Mitchison

**Affiliations:** Max Planck Institute of Molecular Cell Biology and Genetics, Dresden, Germany; Max Planck Institute for the Physics of Complex Systems, Dresden, Germany; Center for Systems Biology Dresden, Dresden, Germany; Cluster of Excellence Physics of Life, TU Dresden, Dresden, Germany; Institute of Science and Technology Austria, Klosterneuburg, Austria; Department of Systems Biology, Harvard Medical School, Boston, United States; Cell Division Group, Marine Biological Laboratory, Woods Hole, United States; Department of Physics, Massachusetts Institute of Technology, Cambridge, United States

## Abstract

Microtubule plus end depolymerization rate is a potentially important target of physiological regulation, but it has been challenging to measure, so its role in spatial organization is poorly understood. Here we apply a method for tracking plus ends based on time difference imaging to measure depolymerization rates in large interphase asters growing in *Xenopus* egg extract. We observed strong spatial regulation of depolymerization rates, which were almost two-fold higher in the aster interior compared to the periphery, and much less regulation of polymerization or catastrophe rates. We interpret these data in terms of a limiting component model, where aster growth results in lower levels of soluble tubulin and MAPs in the interior cytosol compared to that at the periphery. The steady-state polymer fraction of tubulin was ∼30%, so tubulin is not strongly depleted in the aster interior. We propose that the limiting component for microtubule assembly is a MAP that inhibits depolymerization, and that egg asters are tuned to low microtubule density.

## Introduction

Microtubules undergo rapid polymerization dynamics in many cell types, as first revealed by polarization microscopy (Inoué and Sato 1967). Polymerization dynamics occur mostly on plus ends, and are well-approximated by a two-state dynamic model of dynamic instability (Dogterom and Leibler 1993). Plus end behavior is described by four parameters in this model: polymerization, depolymerization, catastrophe and rescue rates. For the robust organization of intracellular structures, these parameters are regulated in both space and time, which also allows for a rapid response to internal and external cues. Most recent studies on the biology and mechanisms of dynamics regulation focused on polymerization and catastrophe rates, in part because multiple factors are known to regulate these rates, and in part because they are the easiest to measure in cells. Tracking of the the plus-tip binding proteins EB1/3 allows measurement of polymerization rate with high reliability, and inference of catastrophe rate from comet lifetime with somewhat lower reliability. In contrast, depolymerization rates are harder to measure. Most studies reporting depolymerization rates focused on pure tubulin systems or on manual measurements of small numbers of microtubules at the periphery of living cells (Desai and Mitchison 1997; Brouhard and Rice 2018). As a result, we know rather little about regulation of depolymerization rates in complex microtubule structures such as the mitotic spindle or microtubule asters. Here, we apply imaging-based methods to systematically measure microtubule depolymerization rates in *Xenopus* egg extract, and report spatial regulation of depolymerization for the first time.

One important emergent property from the four dynamic instability factors, combined with nucleation, is the fraction of polymerized tubulin at steady state, which we will call the “polymer fraction”. In a closed system like a cell, an increase in microtubule mass leads to depletion of soluble subunits which limits further assembly giving rise to a negative feedback of microtubule growth. When continuously supplied with GTP, the system comes to a steady state where the polymer fraction is approximately constant over time, and sufficient tubulin remains in solution to power robust polymerization of GTP-capped plus ends. The steady-state polymer fraction in interphase tissue culture cells was found to be 60-80% (Zhai and Borisy 1994; Kim, Peshkin, and Mitchison 2012). What determines this value, and its signficance for regulation, are largely unknown. Negative feedback from microtubule mass on further assembly in a closed system was conceptualized in the “component limitation” model for mitotic spindle size (Mitchison et al. 2015). A feedback of this kind is inevitable, but its precise mechanism is non-obvious. For pure tubulin, polymerization rate is a linear function of soluble dimer concentration (Desai and Mitchison 1997; Brouhard and Rice 2018). Thus, increased microtubule mass in a closed system decreases polymerization rate by subunit consumption. Catastrophe rate should increase in parallel, assuming it negatively correlates with polymerization rate as in most models (Walker et al. 1988; Gardner et al. 2008). Depolymerization is commonly considered to be a zero-order process whose rate is independent of soluble subunit concentration, though this assumption has been questioned (Gardner et al. 2011). In physiological contexts, tubulin need not be the only limiting component in a closed system. Microtubule-associated proteins (MAPs) that promote microtubule polymerization are also depleted when microtubule mass increases, which could limit further polymerization. It is unclear how MAP-limitation would regulate individual dynamic instability parameters. *Xenopus* egg extract provides undiluted, metabolically-active cytoplasm that reconstitutes physiological polymerization dynamics and is useful for investigating regulatory mechanisms (Field and Mitchison 2018). It allows microscopy-based scoring of large numbers of dynamic microtubules in a relatively homogeneous environment which is ideal for quantitative analysis. Here, we focus on interphase asters, which model the egg-spanning asters that position centrosomes, nuclei and cleavage furrows during early divisions (Wühr et al. 2010). These asters grow to hundreds of microns in radius at a rate of ∼20 microns/min. EB1 tracking showed that the microtubules within them are relatively short, ∼16 microns on average. Their plus end dynamics are in the bounded regime of dynamic instability, so they polymerize transiently, but are biased towards eventual depolymerization with a half-life of ∼1 min (Ishihara, Korolev, and Mitchison 2016). The aster as a whole grows because each microtubule nucleates more than one daughter microtubule during its lifetime by a poorly understood branching process. The density of plus ends is approximately homogeneous within interphase asters, and there are two interesting boundaries: the stationary MTOC and the growing aster periphery. The MTOC nucleates continuously, and is thought to provide the information that directs microtubule polarity. The periphery controls the rate of aster expansion and whether expansion is bounded or unbounded. Plus ends at the periphery polymerize into a microtubule-free environment, while those in the aster interior polymerize into an environment that is dense in microtubules, and as we will show below, partially depleted of soluble subunits. Many questions remain about these asters, including the polymer fraction, minus end behavior, and whether dynamics differ between the interior and periphery. These dynamical properties likely have important implications for aster mechanics and function. Here, we unexpectedly observed that plus end depolymerization rate is subject to strong spatial regulation. We interpret this observation in terms of a new model for regulation of tubulin polymer fraction by MAP depletion.

## Results

### Polymerization rate increases slightly at the aster periphery

We first investigated the spatial variation of plus end polymerization and catastrophe rates using standard EB1 comet tracking methods (Fig. 1A, Movie 1). We found that the polymerization rate remained relatively constant at 30.9 µm/min (N=9348 tracks, SD 5.52, SEM 0.06) as a function of distance from the center of asters (Fig. 1B). We detected a ∼10% increase in polymerization rate at the aster periphery 33.7 ± 10.9 µm/min (mean, std) compared to the interior 30.0 ± 9.5 µm/min (mean, std). The plus end density is lower at the periphery compared to the aster center (Fig. 1C), and resulted in a slight anti-correlation between plus end density and polymerization rate (Fig. 1D). This anti-correlation was much smaller than reported in an independent study (Geisterfer et al. 2020).

**Fig. 1.**
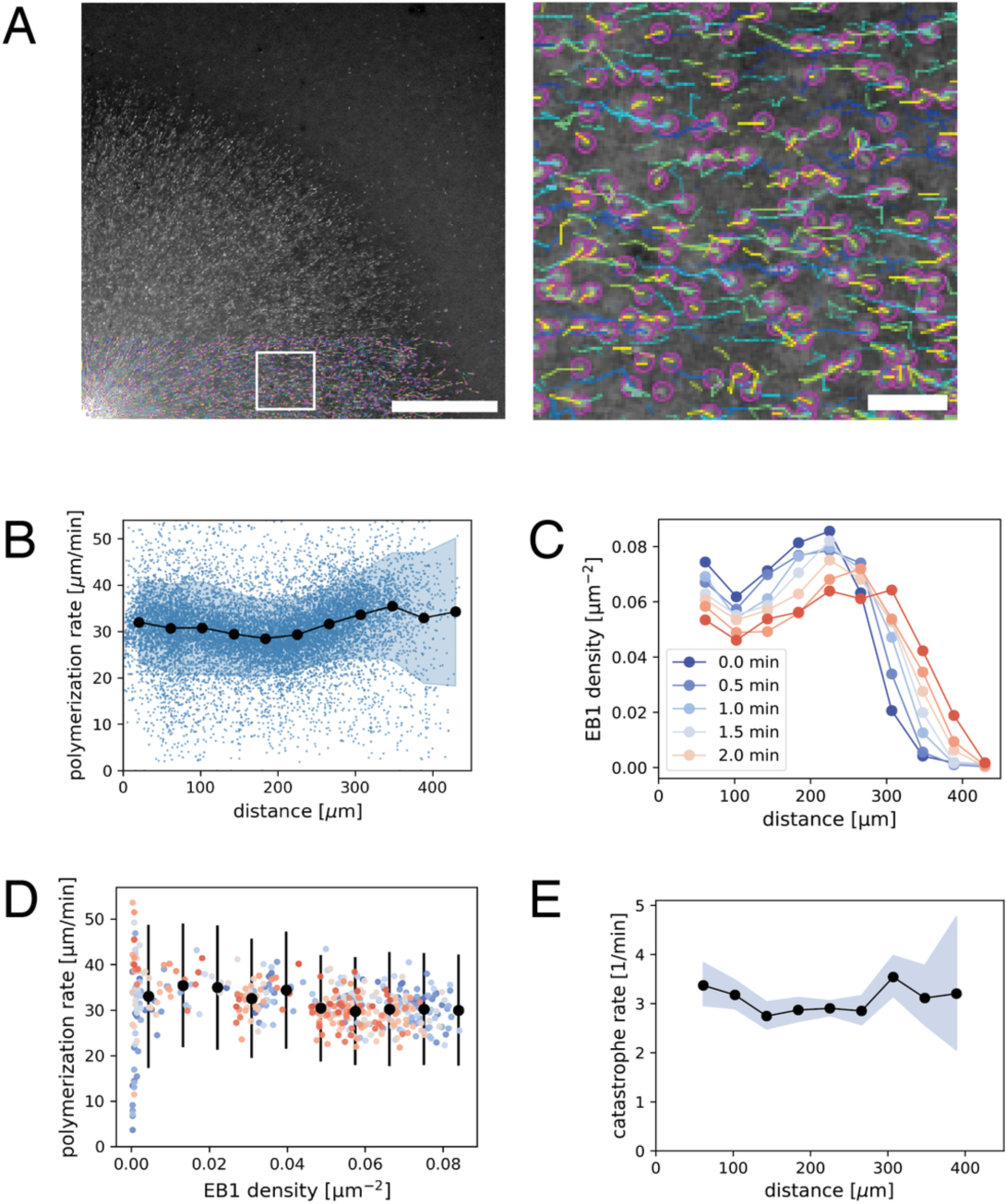
EB1 tracking-based measurements of microtubule plus end polymerization rate and catastrophe rate during aster growth. (A) Time lapse movies of EB1-GFP comets are analyzed by particle tracking. Region in the white box is magnified on the right. Purple spots show the location of individual EB1 comets in this frame, while the trailing lines show the trajectory of the corresponding plus end for the preceding 10 seconds. The velocity of EB1 comets report microtubule polymerization rate. (B) Polymerization rate as a function radial distance from the center of the aster. N=9348 EB1 tracks are represented as blue dots. Filled black circles and the shaded region indicate the mean and the standard deviation of polymerization rate for each spatial bin. The estimated mean difference in the interior (distance>300 µm) vs. periphery (50 µm<distance<280 µm) for t =1.5 min was 3.7 µm/min (95%CI=[2.4, 4.9]). (C) EB1 comet density as a function radial distance. (D) Polymerization rate vs. EB1 comet density. Filled black circles and error bars indicate the mean and the standard deviation of polymerization rate for each EB1 density bin. The different colors correspond to the same time points as in the previous panel. The estimated mean difference in the dense (0.066 1/µm^2) vs. sparse (0.022 1/µm^2) was 3.8 µm/min (95%CI=[2.6, 5.0]). (E) Catastrophe rate is calculated from the duration of EB1 tracks and plotted over radial distance. Filled black circles and the shaded region indicate the mean and the corresponding 95% confidence interval for the catastrophe rate for each spatial bin. Original data used for this analysis is Movie 1. Scale bars 100 µm and 10 µm.

Next, we asked if the plus end catastrophe rate showed spatiotemporal variation using EB1 comet lifetimes as a proxy. The average EB1 comet persisted for 19 seconds which corresponds to a catastrophe rate of 3.1 per minute. This value did not show much variation as a function of location or plus end density (Fig. 1E). These measurements performed with wide-field imaging were consistent with our previous measurements with spinning disc confocal microscopy (Ishihara, Korolev, and Mitchison 2016). Therefore, we expect our measurements to be sensitive to spatial variation in catastrophe rate. In summary, our analysis of EB1 comet imaging demonstrated a modest 10% increase in polymerization rate at the aster periphery, where local plus end density is relatively low, and no measurable differences in catastrophe rate.

### Spatial regulation of depolymerization rates

To measure microtubule depolymerization rates, we applied tubulin intensity difference analysis. This method relies on collecting high quality tubulin images at frequent intervals using TIRF microscopy, subtracting the intensity of sequential frames to generate time difference images, then applying a tracking algorithm. It allows tracking of polymerizing and depolymerizing ends even when most microtubules are in small bundles as is the case in egg extract asters. Our previous analysis of growing and shrinking ends used manual kymograph analysis of difference images and was restricted to the interior of the interphase asters (Ishihara et al. 2014). Here, we applied a newly developed automated workflow (Caldas et al. 2019; Caldas et al. 2020), and systematically compared spatial differences in polymerization dynamics (Fig. 2A). In brief, this method applies a low pass spatial filter to the difference images, followed by particle based tracking to quantify many microtubule ends. Using this workflow, the polymerization rate was found to have a mean of 32.0 ± 4.5 µm/min (mean, std) in the aster interior, and 32.0 ± 4.0 µm/min (mean, std) in the periphery (Fig. 2B, upper panel). These values of polymerization rates are consistent with EB1 comet tracking (Fig. 1). Polymerization rates exhibited a sharp, unimodal histogram, which overlapped for the interior and periphery with no statistical difference. In contrast, the depolymerization rate showed a striking difference in the aster interior vs. periphery. The depolymerization rate was 36.3 ± 7.9 µm/min (mean, std) in the aster interior, compared to 29.2 ± 8.9 µm/min (mean, std) at the aster periphery. The distribution of the depolymerization rates was spread out and had a positive skew in the aster interior, and a negative skew in the aster periphery, with some hint of a bimodal distribution. In summary, our analysis based on an improved intensity difference analysis showed that polymerization rates are similar in the aster periphery relative to the interior, while depolymerization rates show strong spatial variation and are faster inside asters.

**Fig. 2.**
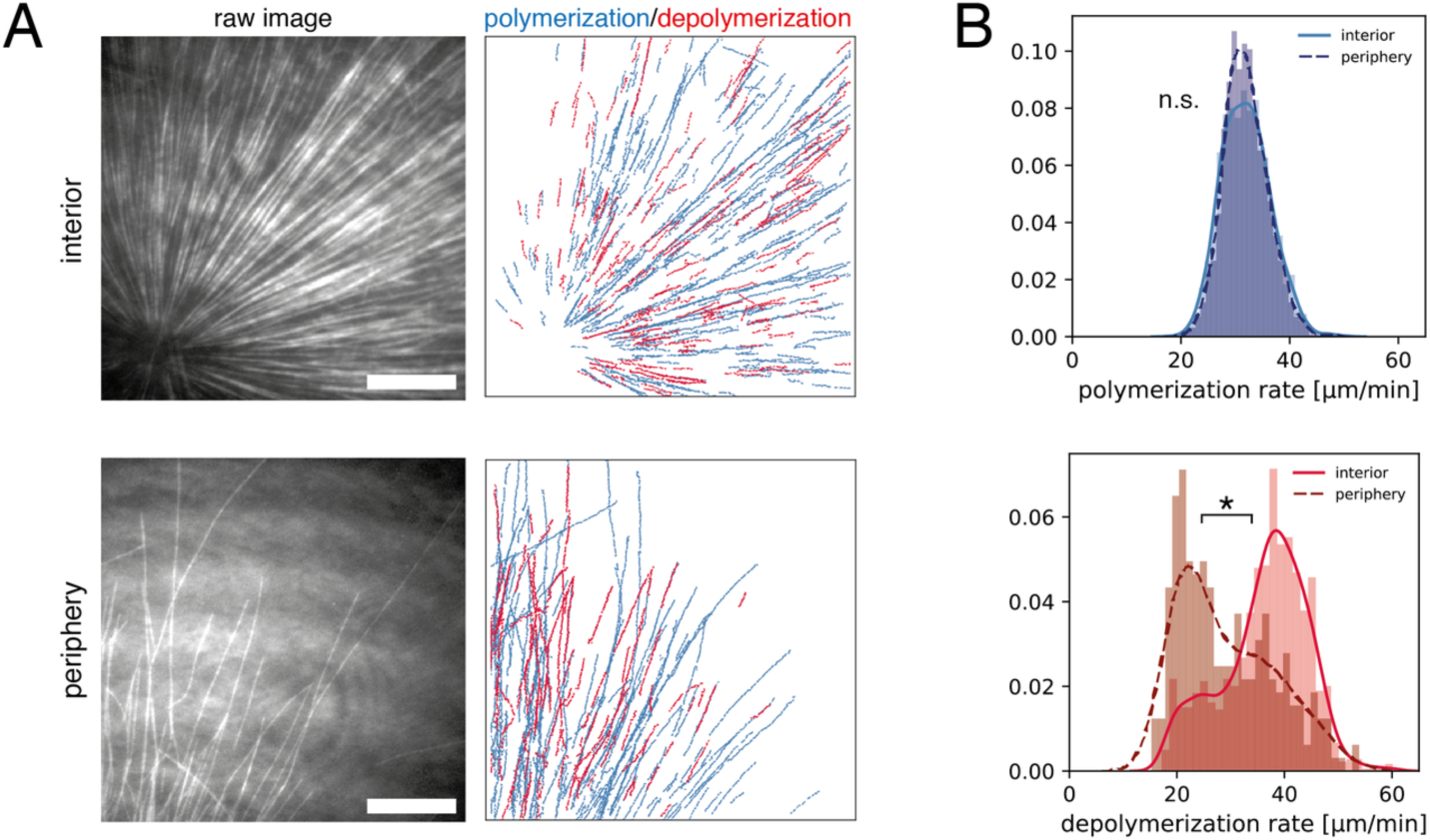
Intensity difference-based measurement of polymerization and depolymerization rates in the interior and the periphery of microtubule asters. (A) TIRF microscopy movies of fluorescent tubulin were subjected to intensity difference analysis, which revealed a spatial map of polymerization and depolymerization for the interior (less than 100 µm away from the MTOC) and peripheral (more than 200 µm from the MTOC with few microtubules) region of microtubule asters. These images correspond to Movies 2 and 3. (B) The distributions of polymerization and depolymerization rates measured from N=6 interior movies and N=6 peripheral movies. The polymerization rate (median 32.0 vs. 32.0 µm/min) showed no statistical difference in the interior vs. periphery (p-value = 0.819, t-test). The depolymerization rate (median 37.6 vs. 26.7 µm/min) was higher in the interior (p-value = 4.44e-16, two sample Kolmogorov-Smirnov test) with estimated difference of mean and median depolymerization rates of 7.1 µm/min (95% CI = [5.9, 8.3]) and 11.0 µm/min (95% CI = [8.4, 12.4]). Scale bar, 20 µm.

Tubulin difference imaging revealed other useful information on polymerization dynamics. Plus end polymerization and depolymerization rates were both variable, but over fairly narrow ranges, so the data supported a two-state model of dynamic instability over alternatives such as a biased random walk (Needleman et al. 2010). Our tracking was not accurate enough to score rate fluctuations within bouts of depolymerization, and we did not try to distinguish stochastic switching between states from more complex dynamics. We observed very few outward-moving depolymerization events, suggesting that minus ends are not depolymerizing, ie there is no significant treadmilling. We also saw very few inward moving polymerization events, suggesting minus ends are static, and perhaps capped.

### Laser ablation induced depolymerization rates

In the conventional model of dynamic instability, growing plus ends lose their structural stability prior to the switch from polymerization to depolymerization. To ask if such physiological catastrophe events are required for the difference in depolymerization rates, we used femtosecond laser ablation to artificially induce depolymerization. Ablation along a line normal to the microtubule axis triggers a spatially defined, synchronous wave of depolymerization (Decker et al. 2018; Decker and Brugués 2015). Using circular patterns, we ablated microtubules at a fixed distance from the center of asters (Fig. 3A, Movie 4). Intensity difference analysis (Fig. 3B) revealed a resulting wave of depolymerization that moved inward at a constant rate (Fig. 3C). This single inward wave is consistent with the fact that (a) the majority of microtubules are oriented with plus ends outward, (b) cut plus ends immediately depolymerize, and (c) newly formed minus ends are stable. By applying this laser ablation assay to multiple asters, we measured an average depolymerization wave velocity of 33.5 ± 9.4 µm/min (mean, std) with a 31.5 µm/min median value, which was comparable to the modal value of the depolymerization rates in the aster interior (Fig. 2). We found no correlation between the rate of depolymerization wave and the distance at which the ablation was induced (15-56 µm). We were not able to reliably cut and measure at the aster periphery due to the low microtubule density. Overall, these observations suggest that depolymerization dynamics are similar for plus ends following a natural catastrophe vs. ablation in the aster interior.

**Fig. 3.**
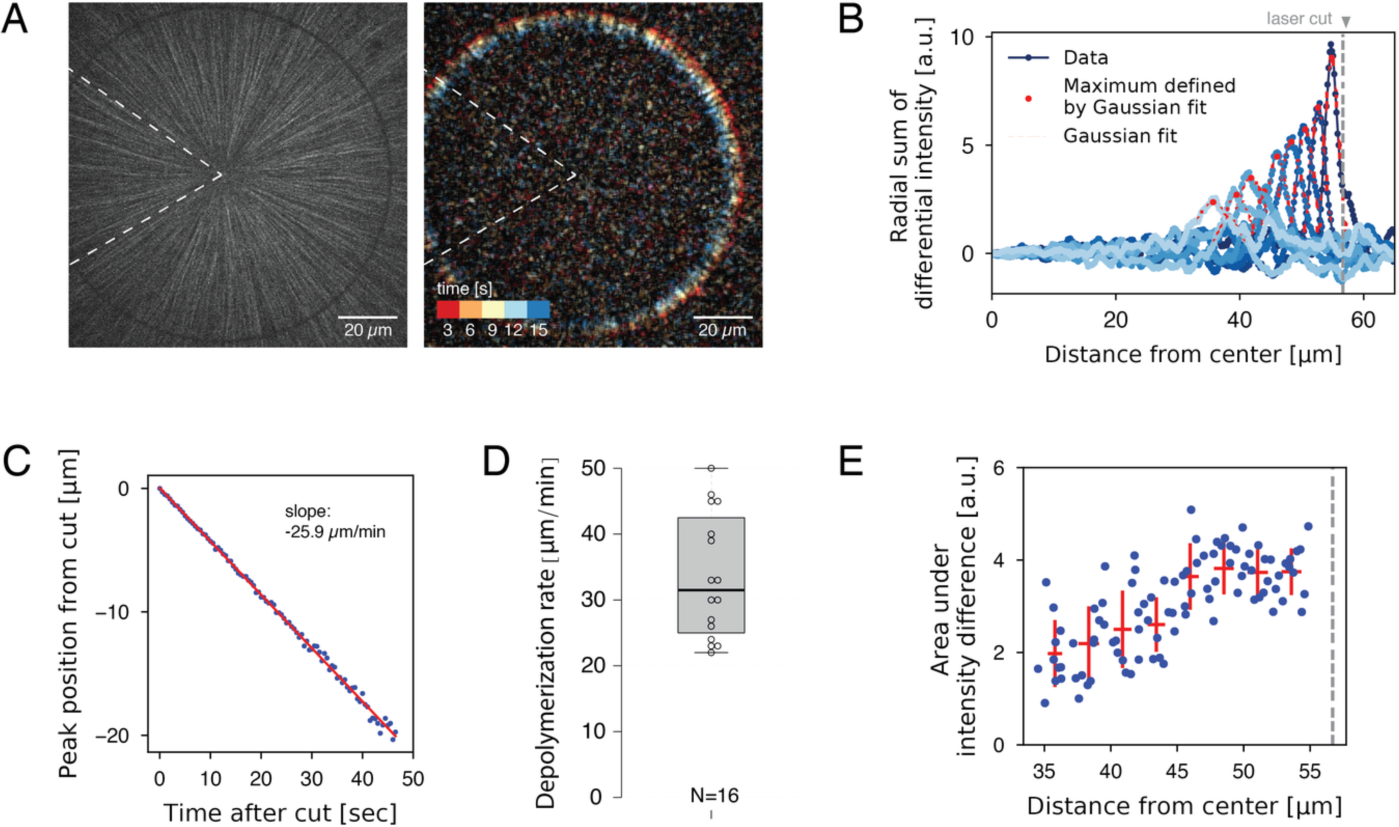
Measurement of depolymerization rates following laser ablation. (A) An interphase aster was subjected to a circular laser cut with radius 56 µm (left), which induced a wave of microtubule depolymerization Movie 4. The resulting movie was used to construct a differential intensity movie (right and Movie 5). Dotted wedge region indicates region excluded from image analysis. (B) Radial sum of differential intensities at different time points (from dark to light blue) of the same laser cut experiment. The area under each curve equals the mass of microtubules depolymerized per time interval of 5 s. Vertical dotted line indicates the location of the laser cut. (C) For each cut, the peak position of the differential intensity travels at constant speed. (D) Depolymerization rates measured from N = 16 laser cuts positioned at 15-56 µm from the center of the aster. (E) The area under differential intensity curves decreases as the depolymerization wave travels inward. This example corresponds to the cut depicted in panel A-C. Vertical dotted line indicates the location of the laser cut.

Laser ablation experiments offer insights into the organization of microtubule structures (Brugués et al. 2012; Decker et al. 2018). We found that the strength of the depolymerization wave (i.e. the area under the curves of the summed differential intensities) decreased with progression (Fig. 3E). This is explained by the fact that depolymerization is halted when individual plus ends reach their corresponding minus ends. Thus, this observation provides evidence that minus ends exist throughout the aster, and confirms our previous proposal that asters are built of short microtubules (Ishihara, Korolev, and Mitchison 2016).

### Aster growth results in spatial gradients of soluble tubulin and EB1

Spatial regulation of depolymerization rate could be due to complex biochemical schemes, such as activation of a dynamics-regulating kinase in the aster interior. We cannot rule out this kind of hypothesis, but we prefer an interpretation based on component limitation, in part because this is a mechanistically simpler hypothesis, and in part because component limitation is inevitable in a closed system. Component limitation is a relevant consideration when comparing microtubule dynamics between the inside and periphery of large asters because aster cytosol is not well mixed by diffusion on relevant time and length scales. Specifically, the aster grows ballistically into fresh cytosol outside the aster much faster than its interior can diffusively equilibrate with that fresh cytosol. A simple Péclet number calculation serves to make this point. Consider the dimensionless Péclet number: 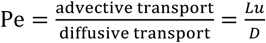. While a typical protein molecule is transported in a diffusive process with cytoplasmic diffusion coefficient 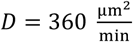 (Salmon et al. 1984), aster growth is a ballistic or advective process, i.e. radius increases linearly with time, with a velocity 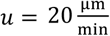 (Ishihara et al. 2014). Given a relevant length scale *L*, we evaluate the magnitude of the Péclet number. The extremes are: Pe>>1, peripheral microtubules grow so fast that they will always grow into fresh cytoplasm with plenty of available material. Pe<<1, aster growth is limited by the diffusive transport of material to the aster periphery. At a depth of *L* = 200 µm from the aster periphery, which is typical for our internal measurements, Pe = 11, so diffusion makes little contribution to supplying subunits. Thus, we expect the aster interior to effectively constitute a closed system, shut off from fresh cytosol supplied by diffusion from the outwardly growing aster boundary.

To seek evidence for depletion of soluble subunits by growing asters we imaged fluorescently labeled tubulin and analyzed its intensity profile as a proxy of local tubulin concentration, summing both polymer and soluble forms (Fig. 4A). The center of asters showed the highest signal, reflecting the highest microtubule polymer density. In addition, we noticed an annular region at the aster periphery with ∼4% lower value of fluorescence signal relative to the background cytosol levels (Fig. 4B). We interpret this as evidence for depletion of soluble tubulin by aster growth. Considering the potential contribution of background signal, the 4% decrease of tubulin intensity is a lower bound for the degree of depletion. The EB1-GFP intensity profile showed a similar spatial pattern, suggesting that microtubule associated proteins may also be depleted to some degree. Finally, we compared the relative position of tubulin/EB1 intensity profiles to that of the EB1 comet density profile (Fig. 4B). This suggested that even the most peripheral plus ends exist in a cytosolic environment that is 4% (or more) depleted of tubulin and EB1 compared to unconsumed cytosol. The half-width of the depleted zone extended ∼50 microns beyond the growing aster periphery, approximately consistent with our Péclet number estimate. This analysis indicated that soluble protein levels may vary within growing asters due to subunit consumption.

**Fig. 4.**
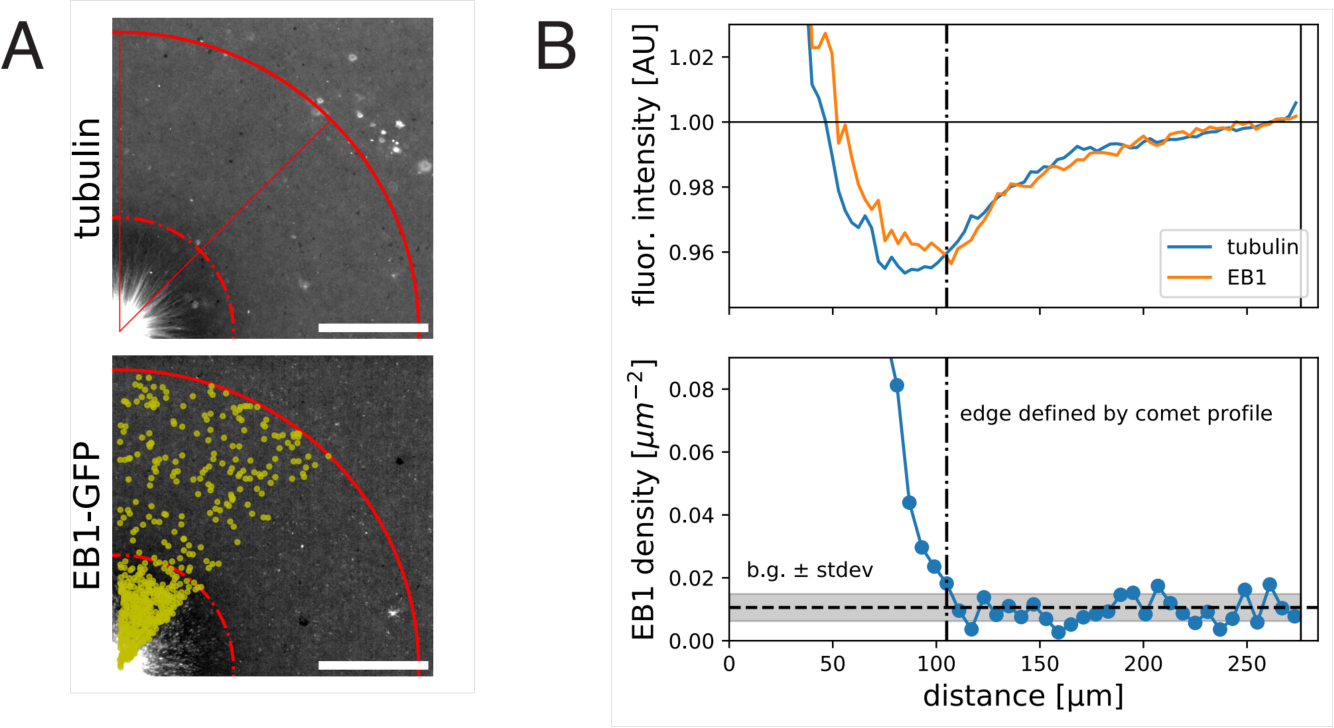
Fluorescent intensity profiles of tubulin and EB1 indicate spatial variation of these species within growing interphase asters. (A) Wide field images of a growing aster visualized with Alexa647-labelled tubulin and EB1-GFP. For both images, the contrast is adjusted to emphasize the zone of low fluorescence intensity. (B) Corresponding quantification of the fluorescence intensity profiles averaged over the quadrant and normalized to the intensity outside the aster (top) and EB1 comet density profile (bottom). We define the aster as the region that has EB1 density higher than the background level + standard deviation. Scale bar, 100 µm.

## Discussion

### Limiting component hypothesis for the spatial regulation of depolymerization rate

The initial motivation of this study was to test our previous assumption that microtubule dynamics was spatially homogenous during aster growth (Ishihara, Korolev, and Mitchison 2016). EB1 tracking experiments showed that the catastrophe rate was largely constant over hundreds of microns (Fig. 1), and the polymerization rate was ∼10% higher in the periphery than the interior. Intensity difference imaging showed that the depolymerization rate varied from modal values of ∼20 µm/min at the aster periphery to ∼40 µm/min in the interior (Fig. 2). Average values were closer, 29 vs. 36 µm/min, but the effect size was evident. This may be the first report of systematic spatial variation in this parameter of dynamic instability.

Multiple hypotheses could be considered to account for the observed spatial regulation of depolymerization rates. We will focus on component depletion models because this is the simplest hypothesis; the growth of structures in confined cytoplasm necessarily consumes the available building blocks and limits further growth (see Introduction). Consistent with such a model, we found evidence for depletion of soluble tubulin and EB1 inside asters by imaging (Fig. 4) and a Péclet number calculation showed that the cytosol inside and outside growing asters do not mix by diffusion once the distance between them is larger than ∼50 microns.

The first candidate for a limiting component in a microtubule system is tubulin itself. As tubulin concentration decreases, the literature predicts that plus ends should grow more slowly and catastrophe more often (see Introduction). However, we observed little spatial variation in these parameters, though EB1-measured growth rates trended in the expected direction (Fig. 1). One reason for a lack of effect of depletion on polymerization rate may be that the steady state polymer fraction in our system is relatively low, ∼30% (Methods, Tubulin polymer fraction estimate), which is the upper bound for difference in soluble tubulin concentration inside and outside asters. Given this small difference and lack of literature on how soluble tubulin levels regulate depolymerization, we explore alternative candidates for the limiting component that is responsible for the spatial regulation of depolymerization rate.

Several MAPs have been shown to slow down depolymerization rates (Drechsel et al. 1992; Andersen and Karsenti 1997). We propose that one or mores MAPs that are present at low concentrations relative to microtubule polymer binds to microtubules at lower density in the interior compared to the periphery, resulting in faster depolymerization in the interior (Fig. 5A). The key parameters that determines whether a given MAP can cause spatial regulation in this way are concentration in cytosol and affinity to the microtubule lattice, which together determine whether the MAP is significantly depleted by microtubule binding. We present a toy model to illustrate how these parameters may help us predict which MAPs regulate depolymerization (see Method, A model of single MAP species binding to microtubule lattice). The model predicts the fraction of freely diffusing MAP molecules 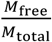 and the site occupancy of the microtubule lattice 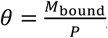, both of which decrease with microtubule polymer concentration P as expected (see Fig. 5B). For a hypothetical MAP whose abundance is 0.2 µM and associates with microtubules with a relatively tight dissociation constant of 100 nM, the fractional occupancy of the microtubule lattice decreases from 0.67 to 0.12 when the polymer concentration increases from 0 to 1.6 µM. We argue that such 5.6-fold difference in microtubule lattice occupancy may be sufficient to cause differences in depolymerization rate that account for spatial regulation. Soluble tubulin, in contrast, only decreases by 0.7-fold. In Table 1, we summarize how different MAPs present in the frog egg are predicted to change their lattice occupancies. Based on the predicted fold change lattice occupancies, we favor EML4, MAP7 and its homologs MAP7D1 and MAP7D2, and MAP1B as candidate regulators of depolymerization that are subject to component limitation effects in large interphase asters, leading to faster depolymerization in the interior.

**Table 1.** Summary of microtubule associated proteins (MAPs) that may slow down plus end depolymerization rate. Protein abundance in the frog egg is from proteome data in (Wühr et al. 2014). For those with reported dissociation constants (Eichenmüller et al. 2001; Andersen and Karsenti 1997; Andersen et al. 1994; Monroy et al. 2018; Tokuraku et al. 2003), we provide the estimated value of θ, the fractional occupancy of the microtubule lattice at low (i.e. excess MAP) and high polymer concentrations (P = 1.6 µM).

| Gene Symbol | Protein Description | abundance [nM] | Kd [nM] | reference | theta at low P | theta at P = 1.6 $\mu\text{M}$ | theta fold change |
| --- | --- | --- | --- | --- | --- | --- | --- |
| MAP4 | Microtubule-associated protein, XMAP230 | 964.5 | 300 | Tokuraku et al. JBC 2003, Andersen et al. JCB 1994 | 0.76 | 0.45 | 1.69 |
| EML2 | Echinoderm microtubule-associated protein-like 2 | 578.18 | ? |  |  |  |  |
| EML4 | EML4 protein | 504.2 | 180 | Eichenmuller et al. Cytoskeleton 2001 | 0.74 | 0.27 | 2.74 |
| EML1 | Echinoderm microtubule-associated protein-like 1 | 135.67 | ? |  |  |  |  |
| MAP7 | Ensconsin | 91.3 | 460 | Monroy et al. Nat. Comm. 2018 | 0.17 | 0.04 | 4.25 |
| MAP1B | Microtubule-associated protein 1B, XMAP310 | 84.68 | 200 | Andersen et al. JCB 1997 | 0.3 | 0.05 | 6.0 |
| MAP1S | MAP1S light chain | 66.04 | ? |  |  |  |  |
| MAP7D1 | MAP7 domain-containing protein 1 | 46.97 | ? |  |  |  |  |
| MAP7D2 | MAP7 domain containing 2 protein variant 2 (Fragment) | 8.55 | ? |  |  |  |  |

**Fig. 5.**
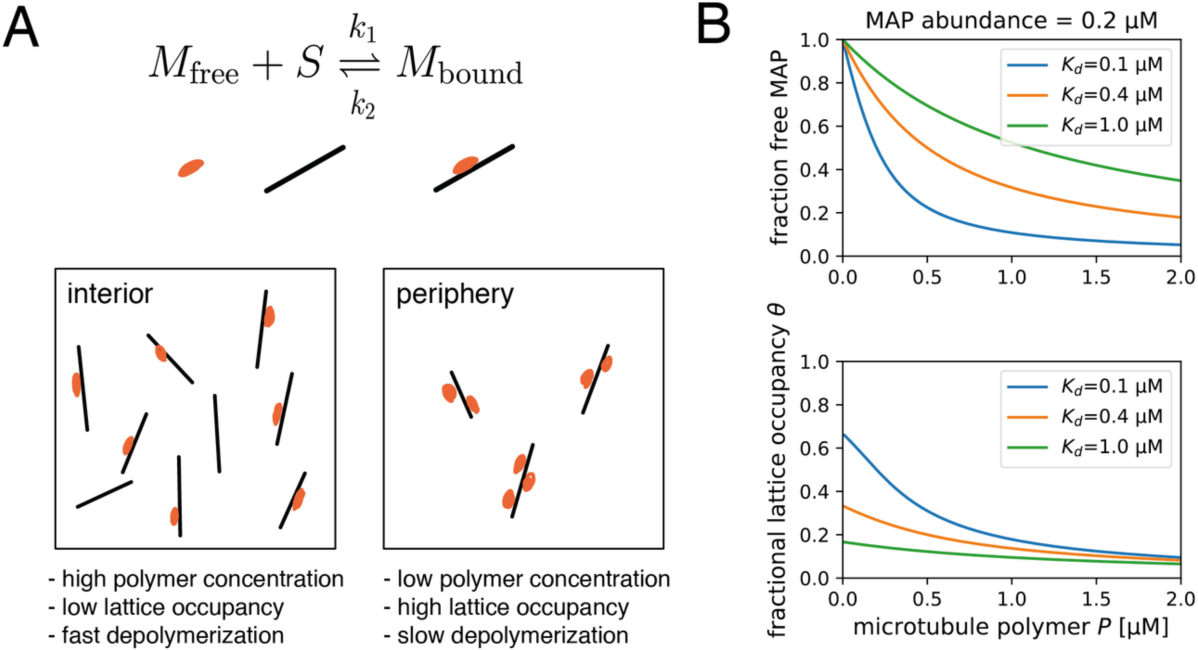
Proposed model for the regulation of depolymerization rate by a MAP species as the limiting component. (A) We consider the equilibrium of a single MAP species that associates with the microtubule lattice with dissociation constant 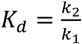. We hypothesize that the degree at which a microtubule is bound by this MAP species slows down depolymerization rate. This effect is greater in the aster periphery, where the microtubule density is lower. (B) Fraction of free MAP and occupied microtubule lattice as a function of microtubule concentration for a hypothetical MAP present at 0.2 µM.

#### Our proposal that a spatially varying MAP

microtubule ratio regulates depolymerization comes with several limitations. First, while a single, low-abundance, tight binding MAP was considered in our model, multiple MAPs may reglated depolymerization rate in eggs. The total abundance of all MAP species easily exceeds 2 µM. Thus, in a more realistic scenario, multiple MAP species may compete for microtubule binding and their order of association is governed by their abundances and affinities. Kinases such as Cdk1, MARK, and Nek may regulate the affinity of these MAPs (Adib et al. 2018; Ookata et al. 1997; Chang et al. 2001; Drewes et al. 1992; Illenberger et al. 1996; Drewes et al. 1997). Our current model is a useful starting point for future studies that consider complex multi-species dynamics of cytoskeletal self-organization including the mitotic spindle (Hazel et al. 2013; Good et al. 2013; Rieckhoff, Ishihara, and Brugués 2019; Rieckhoff et al. 2020).

Second, while most MAPs that slow down depolymerization have been reported to accelerate polymerization in minimal conditions (Drechsel et al. 1992; Andersen et al. 1994; Andersen and Karsenti 1997), we observed little regulation of polymerization rate in our aster reactions. We speculate that, in a cellular context where the synergistic effects of XMAP215 and EB1 play a prominent role in modulating plus end dynamics (Kinoshita et al. 2001; Zanic et al. 2013), our hypothetical MAP may not further accelerate polymerization. It will be interesting to carefully revisit depolymerization rates in other cellular and reconstitution systems.

### Implications of low tubulin polymer fraction for aster mechanics

Finally, what are the implications of our observations for the biology of interphase asters in large eggs? Asters function to separate centrosomes and nuclei at the end of mitosis, and to position cleavage furrows, both of which require that they grow to egg-spanning size. Previously, we modeled aster growth from the interplay of polymerization dynamics and autocatalytic nucleation at the periphery (Ishihara, Korolev, and Mitchison 2016). To prevent the uncontrolled exponential growth expected from an autocatalytic process, we had to assume negative feedback from growth to further assembly, which we implemented as a logistic function that made no mechanistic assumptions. Our current findings suggest that this feedback occurs mainly by increased depolymerization rate in the aster interior, and that it leads to a steady state with a relatively low polymer fraction, estimated as ∼30% tubulin polymerized.

This low polymer density may be important for how dynein exert forces for aster positioning (Wühr et al. 2010). Egg asters are gels comprising a network of relatively short microtubules entangled with F-actin and organelles (Pelletier et al. 2020). Microtubules are the stiffest component of this composite gel, so the bulk stiffness of the aster is likely to depend strongly on the tubulin polymer fraction. We speculate that a low polymer fraction allows the aster gel to deform in response to forces from dynein and the ingressing furrow while maintaining sufficient connectivity to transmit forces across large length scales.

## Materials and Methods

### Aster Reconstitution in Xenopus Egg Extract

Interphase microtubule asters were reconstituted in Xenopus egg extract with anti-AurkA coated beads as microtubule organizing centers (Field, Pelletier, and Mitchison 2017; Field et al. 2014). All reactions were supplemented with 0.04 mg/mL p150-CC1 fragment of dynactin to inhibit dynein motor activity. Fluorescently labelled tubulin or EB1-GFP was used to visualize microtubules.

### Measurement of polymerization and catastrophe rates from EB1 comets

Wide-field images were acquired on a Nikon Eclipse Ni-E upright microscope equipped with a CFI Plan Apochromat Lambda 20× 0.75 N.A. objective lens (Nikon), Nikon motorized XY stage, and Hamamatsu ORCA-Flash4.0 LT scientific CMOS camera, driven by NIS-Elements. EB1-GFP was supplemented to the reactions at a final concentration of 200-400 nM. EB1 comets were imaged at 2 second intervals, and analyzed with TrackMate (Tinevez et al. 2017). For identifying spots, the LoG detector was used with with 2.0 micron spot diameter, threshold 10.0, no median filter, sub-pixel localization enabled. For linking tracks, the LAP tracker was applied with max search radius of 3 micron without gap closing. The results were further analyzed by a custom script written in Python to calculate the polymerization velocity (average of the frame-to-frame velocity for each track) and the catastrophe rate (fitting an exponential function to the comet duration distribution). The DABEST-Python package was used for statistics and effect size estimation (Ho et al. 2019).

### Measurement of polymerization and depolymerization rates of microtubule ends

Physiological microtubule polymerization and depolymerization rates were measured by TIRF microscopy and tubulin intensity difference analysis. Briefly, asters labeled with Alexa647-labelled bovine tubulin were assembled under K-casein coated coverslips and imaged with TIRF microscopy (Ishihara et al. 2014). The Nikon Ti-E motorized inverted microscope was equipped with a Nikon motorized TIRF illuminator, Perfect Focus, a Prior Proscan II motorized stage, Agilent MLC400B laser launch (488 nm, 561 nm, 647 nm), and an Andor DU-897 EM-CCD camera. Alexa 647-labeled bovine tubulin was imaged with a 100× CFI Apo 1.49 N.A. TIRF objective lens (Nikon) with or without 1.5x Optovar. Stream acquisition of images at 500-ms intervals was performed using the RAM capture mode in NIS-Elements software (Nikon Instruments). We defined the aster interior as FOVs with the artificial MTOC in the corner (=center of the FOV less than 100 microns from the center of the aster), while peripheral regions were chosen as the leading edge of the growing aster with very few microtubules in the field of view (=center of the FOV at least 200 microns from the center of the aster).

To obtain spatial information regarding polymerization and depolymerization rates, we applied a recently developed workflow used to quantify treadmilling dynamics in bacterial cytoskeletal filaments (Caldas et al. 2019; Caldas et al. 2020). This method allows to track hundreds of growing and shrinking microtubule ends in an automated fashion, overcoming the limitations of a standard manual kymograph analysis. We first constructed differential time-lapse movies by subtracting the intensity differences between each two consecutive frames. This image-processing step removes static objects from the movie while regions containing intensity fluctuations give rise to either a positive or negative signal (speckles), which corresponds to growth (polymerization) or shrinkage (depolymerization) rates, respectively. Next, the particle-tracking software for ImageJ, TrackMate (Tinevez et al. 2017), was used to automatically detect and follow the resultant moving speckles to rebuild trajectories for analysis. Finally, a custom python script is used to quantify densities and rates of all the detect spots along their trajectories. For TrackMate, we used the LoG (Laplacian Gaussian) detector with an estimated diameter of 1 µm to detect the moving spots. To discard potential false positives, we considered only particles with a signal-to-noise ratio lower than 0.8 and a track displacement distance larger than 1 µm. To build the final trajectories, we used the “Simple LAP tracker” with a “Max Linking Distance” of 1 µm, a “Maximal gap-closing distance” of 1 µm and “Max frame Gap” of 5 frames. Later, we only considered for analysis trajectories longer than 5 sec. To construct the distributions in Fig. 2B, we pooled the frame-to-frame velocities for each track without averaging. The DABEST-Python package was used for statistics and effect size estimation (Ho et al. 2019).

### Measurement of laser cutting-induced depolymerization rates

During the cutting experiments, interphase asters were labeled with Atto565 porcine tubulin and imaged using a Nikon microscope (Ti Eclipse) with Yokogawa CSU-X1 confocal spinning disk, an EMCCD camera (Andor iXon DU-888), a 60x 1.2 NA water immersion objective, and the software AndorIQ for image acquisition. Laser cutting and image analysis were done as described in (Decker et al. 2018) with the only difference that the laser pulse repetition rate was reduced by a pulse picker (APE pulseSelect) from 80 MHz to 20 kHz for some of the cuts, which reduced the probability of destroying the asters. In brief, the sample stage was moved according to the desired cut shape in multiple z planes to reach a cut depth of ∼2 µm around the focal plane. The resulting inward traveling microtubule depolymerization wave was recorded at 2-5 frames/s. The measurement of the polymerization speed involved calculating differential intensities from the raw images with a time difference of 2-3 s (see Fig. 3, Movie 4, 5). These differential intensities were integrated with respect to the radius leading to a Gaussian-shaped integrated intensity profile plotted over the radius for each time point. Using the peak position of fitted Gaussians to these profiles, we determined the distance that the depolymerization wave travelled over time. The slope of this travelled distance over time plot equals the depolymerization speed of the cut microtubules, which was constant over the entire time that the wave was visible (in agreement with depolymerization in metaphase spindles and monopoles (Decker et al. 2018)).

### Fluorescence intensity-based estimation of local depletion of tubulin and EB1

Wide-field images were acquired on a Nikon Eclipse Ni-E upright microscope equipped with a CFI Plan Apochromat Lambda 20× 0.75 N.A. objective lens (Nikon), Nikon motorized XY stage, and Andor Zyla 4.2 Plus scientific CMOS camera, driven by NIS-Elements. Alexa647-labelled bovine tubulin and EB1-GFP were supplemented to the reactions. Dark current subtraction and flat field correction were applied to both channels. A custom script written in Matlab was used to quantify the fluorescent intensity as a function of radial distance from the MTOC (Pelletier et al. 2020). Spot detection of EB1 comets was performed with TrackMate as described above.

### Tubulin polymer fraction estimate

To estimate the fraction of tubulin that is in polymer state, we performed a simple calculation based on our measurements. Specifically, we calculated the total concentration of microtubule polymer in the interior of our aster reactions:

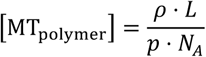

where

*ρ*: number of microtubules per volume [#/µm^3]

*L*: average length of a microtubule – 16 µm (Ishihara 2016)

*p*: pitch of tubulin dimers – 8 nm/13 heterodimers

*N*_*A*_: Avogadro’s number – 6*10^23 heterodimers/mole

By analyzing a z-stack of EB1-GFP images, we measured the thickness of a reaction prepared by squashing 4 µl of extract under a 18×18 mm cover slip as ∼10 micron. Further, we found that a single focal plane of a 20x 0.75 NA lens captures as much as 30-40% of all EB1 comets that are axially distributed in such reaction. Thus, we estimate the microtubule density from 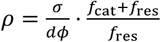.

*σ*: number of microtubule per area in a single focal plane – 0.08 per µm^2

*d*: Thickness of the cover slip reaction – 10 µm

*ϕ*: fraction of EB1 comet detected in a single focal plane - 0.35

*f*_cat_: catastrophe rate - 3.3 [1/min] (Ishihara, Korolev, and Mitchison 2016)

*f*_res:_: rescue rate - 2.0 [1/min] (Ishihara, Korolev, and Mitchison 2016)

The ratio of catastrophe and rescue rates allow us to account for the shrinking plus ends, which are not detected by EB1 comets. We estimate that asters contain [polymer] = 2.65 µM of polymerized tubulin. Comparing this value to abundance of tubulin heterodimers in the frog egg, 8.6 µM (Wühr et al. 2015), we estimate that ∼31 % of tubulin is in the polymer form.

### A model of single MAP species binding to microtubule lattice

We consider a single MAP species M (total concentration *M*_total_) that binds and unbinds from the microtubule lattice (total concentration *P*). Let *S* denote the concentration of free lattice sites and *k*_1_ and *k*_2_ denote the binding and unbinding rates, respectively.

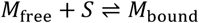

Conservation of MAP species: *M*_total_ = *M*_free_ + *M*_bound_

Conservation of microtubule lattice sites: *P* = *S* + *M*_bound_

The equation for mass action kinetics is:

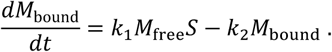

Solving for the steady state solution 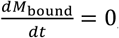, we find:

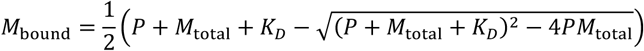

This allows us to calculate, (fraction free M*A*P) 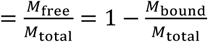 and(fractional lattice occupancy)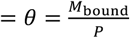.

## Supplementary Material

**Movie 1**. Timelapse imaging of EB1-GFP comets during aster growth.

**Movie 2**. TIRF microscopy-based intensity difference imaging of the aster interior. Blue correspond to positive intensity difference interpreted as polymerization, while red correspond to negative intensity difference interpreted as depolymerization.

**Movie 3**. TIRF microscopy-based intensity difference imaging of the aster periphery.

**Movie 4**. A circular laser ablation induces an inward wave of depolymerization.

**Movie 5**. Intensity difference movie corresponding to the laser ablation in Movie 4.

## Author Contributions

KI and TJM designed the research and wrote the manuscript with help from all authors. TJM acquired the data and KI performed the analysis for EB1 comets. KI, PC, and ML performed the tubulin intensity difference analysis. KI, FD, and JFB performed the laser ablation experiments and analyzed the data. KI and JFP performed the analysis to infer local protein depletion from wide field microscopy images. KI developed the theoretical model.

## Acknowledgements

The authors thank the members of Mitchison, Brugues, and Jay Gatlin groups (Uni. of Wyoming) for discussions. We thank Heino Andreas (MPI-CBG) for frog maintenance. We thank Nikon Inc. for microscopy support at MBL. KI was supported by fellowships from the Honjo International Scholarship Foundation and Center of Systems Biology Dresden. FD was supported by the DIGGS-BB fellowship provided by the DFG. PC is supported by a Boehringer Ingelheim Fonds (BIF) PhD fellowship. JFP was supported by a fellowship from the Fannie and John Hertz Foundation. ML’s research is supported by a European Research Council (ERC) grant ERC-2015-StG-679239. JB’s research is supported by the Human Frontiers Science Program (CDA00074/2014). TJM’s research is supported by NIH grant GM39565 and by MBL summer fellowships.

